# Antiviral activity of pacific oyster (*Crassostrea gigas*) hemolymph against a human coronavirus

**DOI:** 10.1101/2021.11.01.466695

**Authors:** Rebecca L. Pedler, James O. Harris, Peter G. Speck

## Abstract

Coronaviruses can cause severe respiratory infections in humans. In this study we assessed the antiviral activity of Pacific oyster (*Crassostrea gigas*) hemolymph against a human coronavirus, HCoV-229E. An eight-fold reduction in infectivity of HCoV-229E on Huh-7 cells was observed in the presence of 10% *C. gigas* hemolymph. Antiviral activity of *C. gigas* hemolymph positively correlated with its concentration and appears to be active during an intracellular stage of HCoV-229E infection.

---

Human coronaviruses are enveloped, single stranded RNA viruses that are further classified as alpha-coronaviruses (human coronavirus-229E (HCoV-229E) and HCoV-NL63) or beta-coronaviruses (HCoV-OC43, HCoV-HKU1, middle eastern respiratory syndrome (MERS-CoV) and severe acute respiratory syndrome (SARS-CoV and SARS-CoV-2) (1). SARS-CoV-2 is a novel human coronavirus which emerged in December 2019 as the causative agent of coronavirus disease 2019 (Covid-19) (2-4). Safe and effective antiviral treatments for SARS-CoV-2 are yet to be identified, despite multiple drug repurposing attempts (5-7).

Marine molluscs represent an unexploited source of medicinal compounds (R.L. Pedler, and P.G. Speck, submitted for publication) (8-11). Marine molluscs lack an adaptive immune system and exclusively elicit innate immune responses (12-14), while living in an environment containing virus particles in the order of >10^7^ per ml (15, 16). This demonstrates the success of their strategies to prevent viral infection, which includes production of potent antiviral compounds (11, 13). *In vitro* inhibition of HSV-1 has been observed using extracts from the common cockle (*Cerastoderma edule*), greenlip abalone (*Haliotis laevigata*) (17), Japanese carpet shell (*Ruditapes philippinarum*), European flat oyster (*Ostrea edulis*), common whelk (*Buccinum undatum*) (18), blacklip abalone (*Haliotis rubra*) (19, 20), veined rapa whelk (*Rapanosa venosa*) (21) and Mediterranean mussel (*Mytilus galloprovincialis*) (22). Extract from the flesh of the red abalone (*Haliotis rufescens*) protects mice against poliovirus and influenza A (23, 24), while paolin II from the Eastern oyster (*Crassostrea virginica*) inhibits poliovirus (25).

Hemolymph of the Pacific oyster (*Crassostrea gigas)* has *in vitro* antiviral activity against HSV-1 and adenovirus respiratory strain 5 (AdV-5) (26-28). The major *C. gigas* hemolymph protein, cavortin, exerts an antiviral effect against HSV-1 after entry into Vero cells (26). Cavortin is a Mr 20,000 protein which acts as a metal chaperone (29). Intracellular zinc has therapeutic potential for SARS-CoV-2 (30-32), and its efficacy is improved by coupling with a metal chaperone (31, 33). *Crassostrea gigas* has a high zinc content (34) and given cavortin’s suggested role as a metal chaperone (29), it is possible that *C. gigas* cavortin has potential antiviral activity against SARS-CoV-2 and may facilitate zinc transport into host cells.

The discovery of antiviral agents for SARS-CoV-2 is challenged by the limited number of laboratories with the appropriate biosafety containment level (35, 36). HCoV-229E can be handled in lesser-rated laboratories making it more accessible for research on human coronaviruses (37) and this virus could be used for initial screening for anti-coronavirus activity. This study is the first, that we are aware of, which assesses antiviral activity of *C. gigas* hemolymph against HCoV-229E.

Huh-7 cells, obtained from M. Beard (Adelaide University, South Australia), were grown in Dulbecco’s modified eagle medium (DMEM) (Gibco #11965118) supplemented with 10% foetal bovine serum (FBS) (Gibco #10099141), according to standard methods (38). Twelve *C. gigas* oysters, grown in Coffin Bay, South Australia, were purchased from local seafood merchants. After opening, *C. gigas* hemolymph was extracted from the pericardial cavity using a sterile syringe and 27g needle. Hemolymph was pooled, filter sterilised using a 0.2*μ*m filter and stored at -20°C until required. Cytotoxicity of *C. gigas* hemolymph was determined using a trypan blue exclusion assay (26). Huh-7 cells were seeded into a 24-well plate with medium as above and 0%, 2%, 5%, 10% or 20% (v/v) *C. gigas* hemolymph. Cells were incubated for two days at 37°C in 5% CO_2_, before being stained *in situ* with 0.4% trypan blue (Gibco #15250061) (26). The number of non-viable cells in three different fields of view were then counted under an Olympus CK2 microscope at 40x magnification (39). The mean number of non-viable cells was lowest for 0% (1.00 ± 0.00 cells) and 2% (1.00 ± 0.47 cells) (Fig. 1). Huh-7 cell death sharply increased as hemolymph concentration exceeded 10% (Fig. 1), therefore, 10% was considered an appropriate concentration for use in anti-HCoV-229E assays. In Vero cells, *C. gigas* hemolymph can cause 10% cell death at a concentration of 13% (v/v) (26) or 50% cell death at 750µg ml^-1^ (27). The acellular and cellular fractions of *C. gigas* hemolymph can cause 50% cell death in Hep-2 cells at concentrations of 0.32mg ml^-1^ and 0.19mg ml^-1^ respectively (28).

**FIG. 1.**
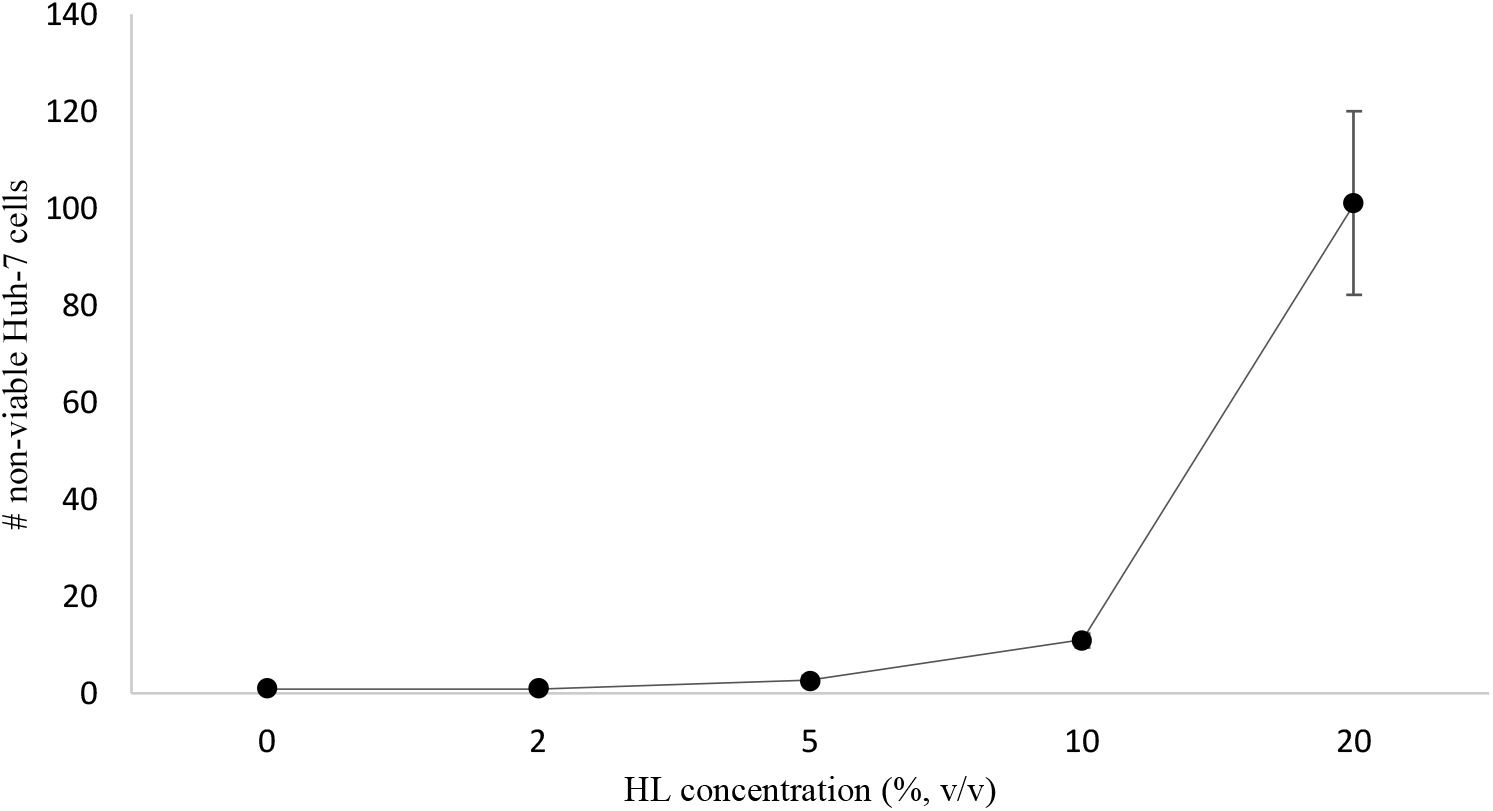
Mean (± standard deviation) number of non-viable Huh-7 cells treated with varying concentrations of Pacific oyster (*Crassostrea gigas*) hemolymph (0, 2, 5, 10, 20% v/v).

**FIG. 2.**
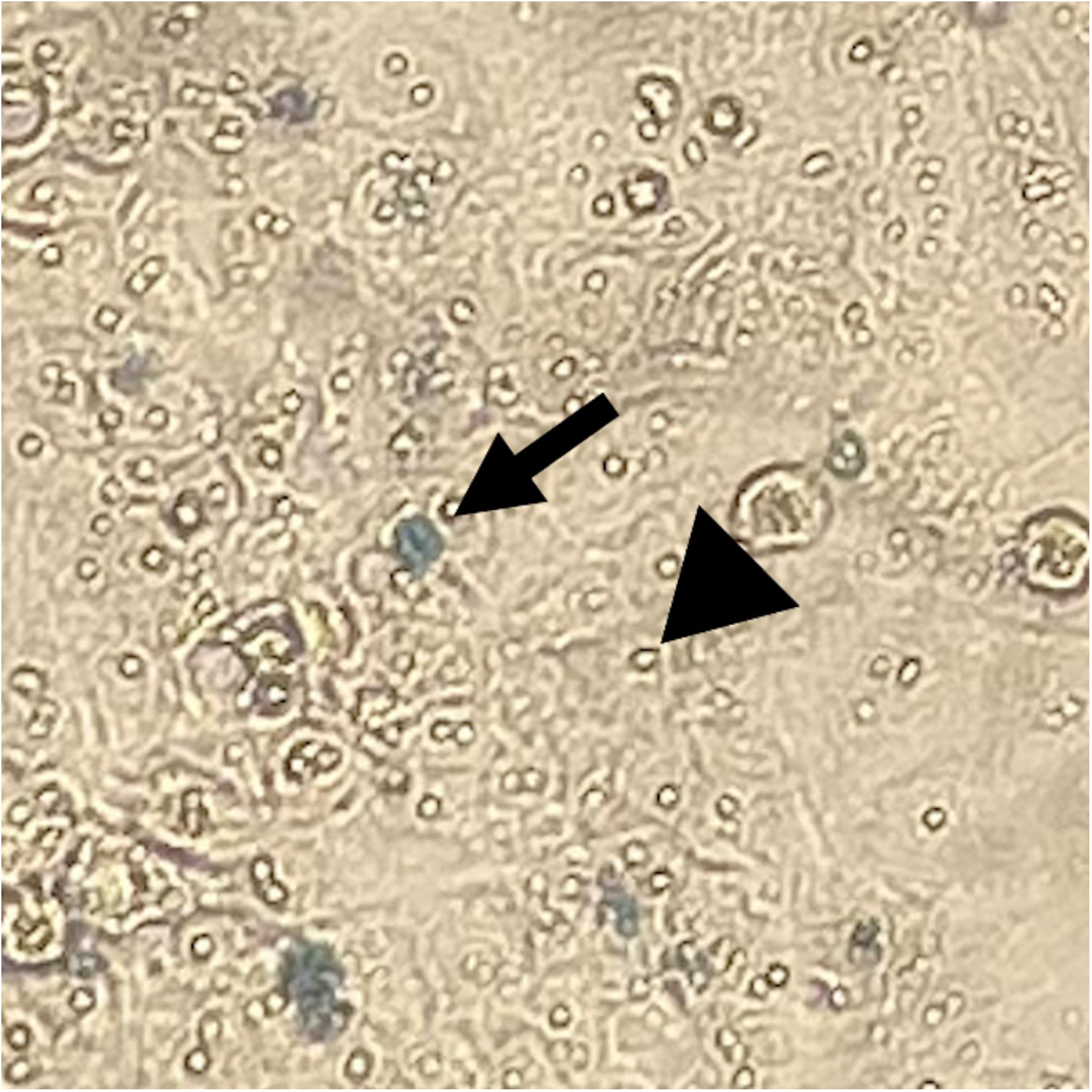
non-viable (arrow) and viable (arrowhead) Huh-7 cells treated with 20% Pacific oyster (*Crassostrea gigas*) hemolymph.

HCoV-229E was obtained from H. Whiley (Flinders Water Quality and Health Research Consortium, South Australia). Virus titres were determined as 50% tissue culture infective doses (TCID_50_) (40). Huh-7 cells were seeded into 96-well plates and either 0 or 10% *C. gigas* hemolymph. Three ten-fold dilutions, followed by eight two-fold dilutions were prepared using HCoV-229E stock and DMEM and inoculated into 96-well plates. Cells were incubated at 37°C in 5% CO_2_ for five days before being fixed with 10% formaldehyde and stained with 0.5% crystal violet (Thermo #S25275B). Wells illustrating cytopathic effect were counted, allowing TCID_50_ to be calculated using the Reed-Muench method (41). When Huh-7 cells were assayed with 10% *C. gigas* hemolymph, an eight-fold reduction in the HCoV-229E titre (4.00 × 10^5^ TCID_50_ ml^-1^) and an antiviral activity of 87.5% was observed (Table 1). A dose-response curve was generated using 0%, 2%, 5%, 10% and 15% *C. gigas* hemolymph which revealed that antiviral activity positively correlated with its concentration (Table 1, Fig. 3). This is consistent with the dose-dependent antiviral activity of *C. gigas* hemolymph protein, cavortin, against HSV-1 (26).

**TABLE 1.** Virus titre values (TCID_50_ ml^-1^) for human coronavirus 229E (HCoV-229E) in Huh-7 cells treated with varying concentrations (0, 2, 5, 10, 15% v/v) of Pacific oyster (*Crassostrea gigas*) hemolymph.

| Extract/control | Virus titre (TCID<sub>50</sub> ml<sup>-1</sup>) | Antiviral activity: %<br>Reduction in virus titre |
| --- | --- | --- |
| Negative control (DMEM) | $3.20 \times 10^6$ | 0.00 |
| 2% *C. gigas* hemolymph | $8.00 \times 10^5$ | 75.00 |
| 5% *C. gigas* hemolymph | $7.92 \times 10^5$ | 75.24 |
| 10% *C. gigas* hemolymph | $4.00 \times 10^5$ | 87.50 |
| 15% *C. gigas* hemolymph | $2.83 \times 10^5$ | 91.16 |

**FIG. 3.**
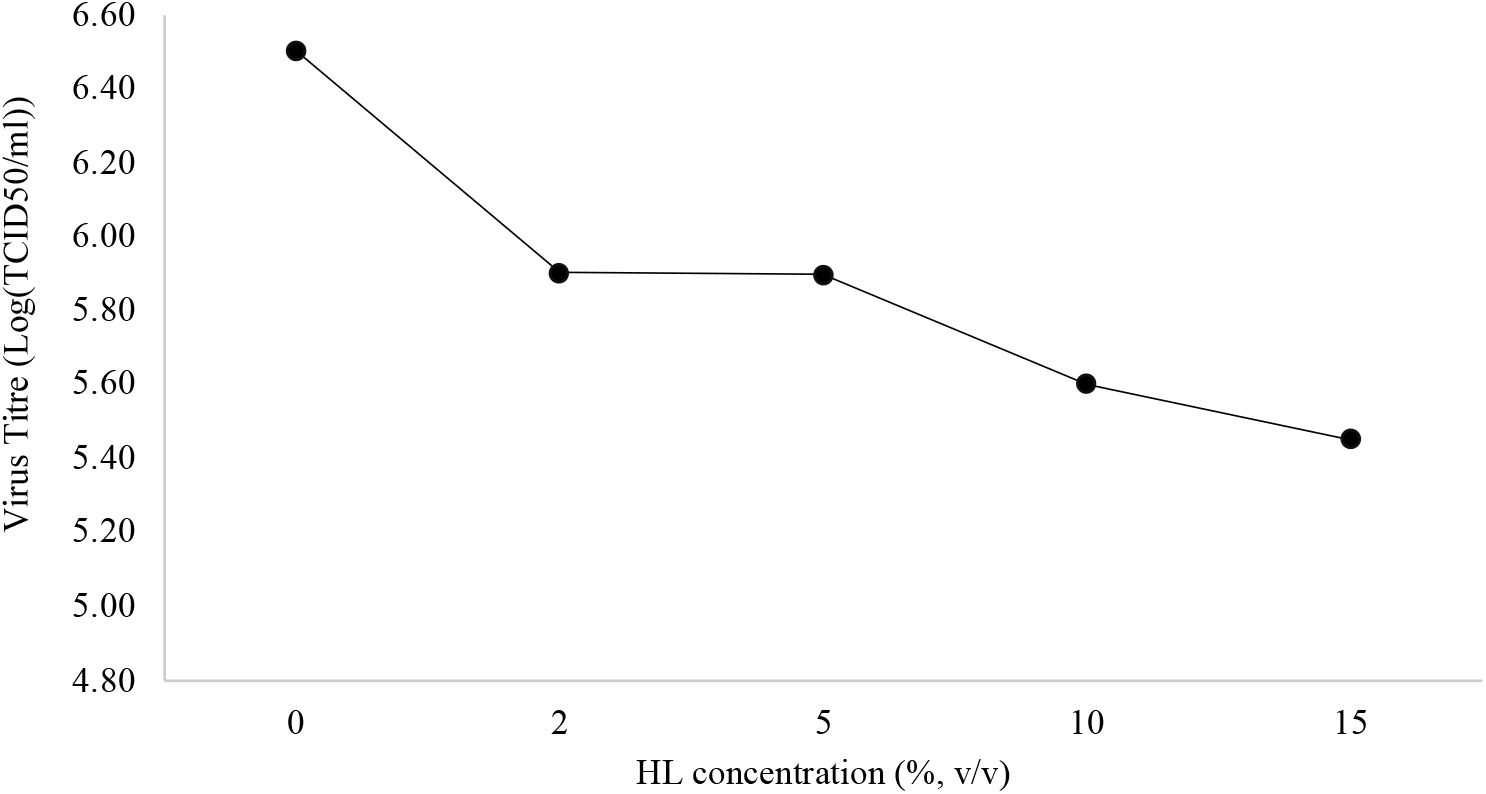
Virus titre values (Log(TCID_50_ ml^-1^)) for human coronavirus 229E (HCoV-229E) in Huh-7 cells treated with varying concentrations (0, 2, 5, 10, 15% v/v) of Pacific oyster (*Crassostrea gigas*) hemolymph (HL).

Time of addition assays were used to determine the stage of HCoV-229E infection targeted by *C. gigas* hemolymph. In previous studies, the greatest antiviral protection of Vero cells from HSV-1 infection was observed when *C. gigas* hemolymph was added 0-2 hours after infection (26, 27), suggesting that the antiviral effect was most likely exerted after virus attachment and entry. An intracellular mode of antiviral action has been observed for other molluscan compounds, including lipophilic extract of *H. laevigata* (17) and myticin C peptides from *M. galloprovincialis* (22). Here, Huh-7 cells were seeded into 24-well plates with DMEM and 10% FBS. A series of dilutions (10^−4^, 10^−5^, 10^−6^ and 10^−7^) were prepared using HCoV-229E stock and DMEM and inoculated into plates. *C. gigas* hemolymph was added either immediately or 60 minutes after HCoV-229E and a negative control plate was also prepared. There was little difference in *C. gigas* hemolymph antiviral activity when it was added to Huh-7 cells immediately (98.21%) or 60 minutes after HCoV-229E infection (96.11%) (Table 2). This suggests that *C. gigas* hemolymph most likely acted during an intracellular stage of HCoV-229E infection. Antiviral compounds which act during an intracellular stage of HCoV-229E infection have been identified (42-44). The macrolide and immunosuppressive compound, FK06 inhibits HCoV-229E replication in Huh-7 cells (44), while the antimalarial drug chloroquine inhibits HCoV-229E replication in epithelial lung cells (L132) by suppressing P38MAPK (43). Thapsigargin, from the Thapsia (*Thapsia garganica*) plant acts during an intracellular stage of HCoV-229E infection either by inhibiting replication or activating unknown antiviral effector systems in Huh-7 cells (42).

**TABLE 2.** Virus titres (TCID_50_ ml^-1^) and antiviral activity (% reduction in virus titre) for human coronavirus 229E (HCoV-229E) in Huh-7 cells treated with either 0% (negative control) or 10% Pacific oyster (*Crassostrea gigas*) hemolymph.

| Extract/control | Time of extract addition | Virus titre (TCID<sub>50</sub> ml<sup>-1</sup>) | Antiviral activity: % reduction in virus titre |
| --- | --- | --- | --- |
| Negative control (DMEM) | - | $2.79 \times 10^8$ | 0.00 |
| 10% *C. gigas* hemolymph | Immediately after infection | $5.00 \times 10^6$ | 98.21 |
| 10% *C. gigas* hemolymph | 60minutes after infection | $1.08 \times 10^7$ | 96.11 |

This study reveals that *C. gigas* hemolymph has *in vitro* antiviral activity against human coronavirus HCoV-229E. This finding is relevant in the current pandemic and reinforces that *C. gigas* hemolymph has broad-spectrum antiviral activity. Further research is required to identify and characterise the antiviral compound(s) produced by *C. gigas*.

